# A short commentary on indents and edges of β-sheets

**DOI:** 10.1101/850982

**Authors:** Harshavardhan Khare, Suryanarayanarao Ramakumar

## Abstract

β-sheets in proteins are formed by extended polypeptide chains, called β-strands. While there is a general consensus on two types of β-strands, viz. ‘edge strands’ (or ‘edges’) and ‘inner strands’ (or ‘central strands’), the possibility of distinguishing between different regions of inner strands remains less explored. In this paper, we address the portions of inner strands of β-sheets that stick out on either or both sides. We call these portions the ‘indent strands’ or ‘indents’ because they give the typical indented appearance to β-sheets. Similar to the edge strands, the indent strands also have β-bridge partner residues on one side while the other side is still open for backbone hydrogen bonds. Despite this similarity, the indent strands differ from the edge strands in terms of various properties such as β-bulges and amino acid composition due to their localization within β-sheets and therefore within folded proteins to certain extent. The localization of indents and edges within folded proteins seems to govern the strategies deployed to deter unhindered β-sheet propagation through β-strand stacking interactions. Our findings suggest that, edges and indents differ in their strategies to avoid further β-strand stacking. Short length itself is a good strategy to avoid stacking and a majority of indents are two residue or shorter in length. Edge strands on the other hand are overall longer. While long edges are known to use various negative design strategies like β-bulges, prolines, strategically placed charges, inward-pointing charged side chains and loop coverage to avoid further β-strand stacking, long indents seem to favor mechanisms such as enrichment in flexible residues with high solvation potential and depletion in hydrophobic residues in response to their less solvent exposed nature. Such subtle differences between indents and edges could be leveraged for designing novel β-sheet architectures.

## Introduction

β-sheets are one of the fundamental secondary structural elements of proteins and form important structural scaffolds. They consist of many β-strands arranged adjacent to each other in parallel, antiparallel or mixed orientations and are held by periodic main chain hydrogen bonds along with other potential forces such as van der Waals interactions, electrostatic interactions, ionic bonds and hydrophobic effects. The immunoglobulin fold is one classic example of robust β-sheet scaffold holding the hypervariable loops in position. β-barrels are also a good example of structurally robust β-sheet architectures.

Another well studied example involving β-sheets is amyloid fibrils [1]. Amyloid fibrils are perhaps one of the most stable protein structures. Unhindered stacking of β-strands results in the formation of amyloid fibrils. Different sequences can give rise to amyloid fibrils composed of β-strands assembled in a long β-sheet structure. Recent studies have shown that while the governing principles remain same, the amyloid fibrils exhibit polymorphism [2]. Different types of amyloid fibrils are considered to be one of the causes of protein aggregation diseases such as Alzheimer’s disease, Parkinson’s disease, Type-II diabetes and ALS (Amyotrophic Lateral Sclerosis).

Exceptions to the β-sheets in either globular proteins or amyloids exist as isolated extended β-strands and have been studied comprehensively by Eswar et al. [3].

While β-sheets exhibit robust architectural properties in many proteins, their existence is not limited to being only structural scaffolds. β-sheets are also important in functions of proteins, for example, ligand binding or target recognition [4], DNA binding [5, 6], protein-protein interactions (PPIs) [7] and targets of proteases [8].

Importance of β-sheets and β-strands in PPIs has been known for a long time. Remaut and Waksman [7] have summarized PPIs through various ways of addition of β-strands to existing β-sheets. Analysis of protein-protein interfaces by Watkins and Arora [9] highlights importance of β-strands as useful targets of modulation of PPIs. β-strand–β-strand associations have also been leveraged to design peptidomimetic ligands that inhibit PPIs [10].

Besides PPIs in globular proteins, amyloid fibrils also propagate through β-strand stacking interactions. Various approaches for modulation of amyloid aggregation have been explored, for example, inhibitors [11], covalent modulators [12], non-covalent modulators [13], stabilizers of monomer Aβand destabilizing agents of pre-formed fibrils [14, 15], β-breaker substituted endomorphin analogue [16] and self-inhibition using fragments of an amyloid peptide [17, 18].

Despite such efforts, designing β-sheets, β-strands and their ligands still remains more challenging than that for any other protein secondary structural element. It is thus important to investigate deeper into the fundamental aspects of β-sheets.

Unhindered stacking of β-strands is not exhibited by majority of the natural β-sheets. Many β-sheets have progressively shorter strands towards edges thereby terminating β-sheet propagation. Most β-sheets terminate in two strands, commonly called ‘edges’ or ‘edge strands’. The remaining strands are called ‘inner strands’. Short length of edges is one of the negative design strategies [19] employed by β-sheets to deter further β-strand stacking. Several examples exist where edges are considerably long and have been shown to exhibit different negative design principles to avoid β-strand stacking, such as inward pointing charged side-chains, β-bulges and loop coverage [19].

The shortening of β-strands towards edges of β-sheets gives rise to edge-like flanking regions of inner strands. In most studies on β-sheets, these regions have not been differentiated from inner strands [20, 21, 22, 23, 24]. In fact, these regions are similar to the terminal edge strands in having only one β-strand partner. However, their localization in the β-sheets, and therefore in the folded proteins, differs from that of the edge strands. In this paper, we treat these edge-like flanking regions of inner strands as different type of strands. We call these regions ‘indents’ or ‘indent strands’ to reflect their property to give an indented appearance to β-sheets (Figure 1). Zhang et al. [25] have provided a good statistical analysis of length distribution of indents which they call the ‘terminal extensions’. However, to the best of our knowledge, the comparison of indents and edges with respect to amino acid residue preferences still remains unexplored. The main purpose of this work is to investigate whether edges of β-sheets are different from indents in terms of amino acid residue composition and the strategies they employ to deter further β-strand stacking. Understanding these differences might be useful in *de novo* design of β-sheet structures.

**Figure 1:**
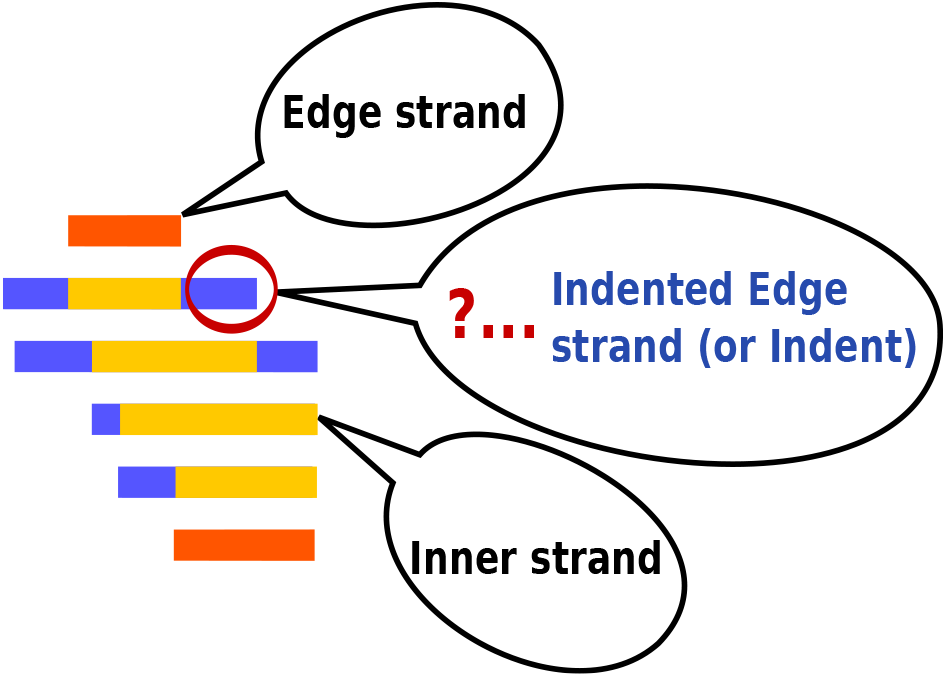
Types of β-strands

## Materials and Methods

### Dataset generation

This analysis was performed on X-ray crystal structures obtained from the PDB (Protein Data Bank). Structures determined using NMR and modeled structures were not included in this analysis. A dataset of 11403 high resolution (< =2.0 Å) protein structures from the PDB was culled using PISCES culling server [26] with R-factor < = 25% and sequence identity < = 25%. Out of 5827 chains obtained after culling, 4415 contained at least one β-sheet (identified using the DSSP program [27]). This set of 4415 chains was used for further analyses. β-bulges were identified using PROMOTIF 3.0 program [28]. Solvent accessibilities were calculated using NACCESS program [29]. Composition profiler [30] was used to calculate enrichment and depletion of various amino acid residues and physico-chemical properties. In-house python scripts were written analyze the protein structures and identify various β-strand types.

### 2D-map representation of β-sheet

A 2D-map (2 Dimensional map) representation of β-sheet architecture has been used (Figure 2) to identify indent strands. The residues in the β-sheet form β-bridge pairs on either sides across β-strands. These pairings were used to identify hypothetical β-bridge sequences that run across the β-sheet. These β-bridge sequences terminate either in an edge residue or an indent residue. This information was used to identify portions of inner strands that stick out to form indents. In-house python scripts were written to transform the β-sheet structures obtained from the 3D coordinates in the PDB files to the 2D-map representation and to identify amino acid residues in edge strands, indent strands and interior of the β-sheet for further analysis.

**Figure 2:**
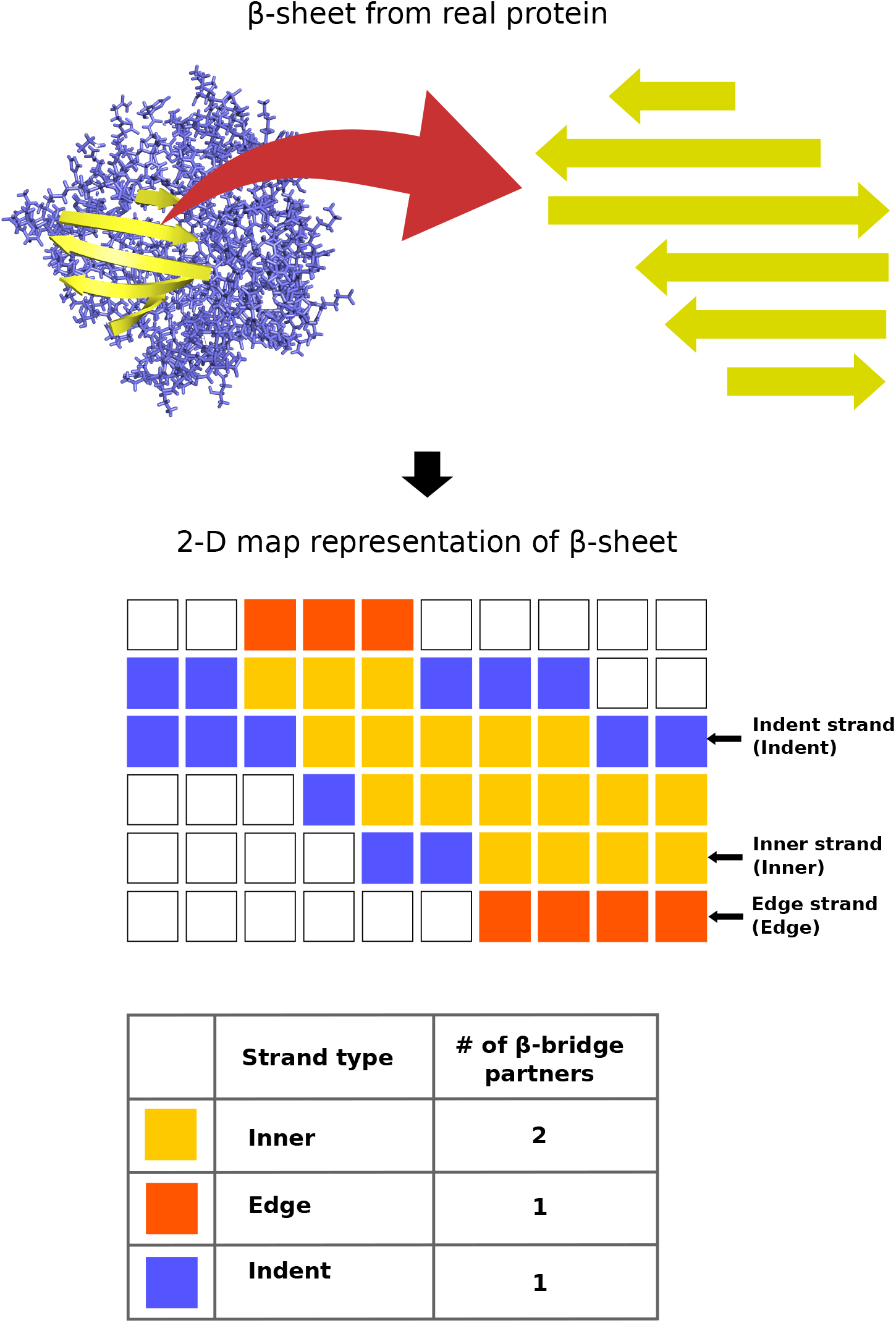
2D-map representation of β-sheet

## Results and Discussion

### Basic statistics of the data

From the 4415 protein chains that were analyzed, 7079 β-sheets were obtained which contained a total of 43676 β-strands ranging between 1-8 residue length. The distribution of these β-strands in the range of 1-8 residue length shows that approximately 38 % (i.e. 16749) of the β-strands are edge strands (Table 1). The remaining 62 % (i.e. 26927) inner strands were analyzed further to identify the indents. These 26927 inner strands contained 37999 indents, which means on an average ≈1.4 indents per inner strand, suggesting at least one indent in each inner strand (maximum can be two, one on each side). The indents thus seem to be abundant in β-sheets.

**Table 1:**
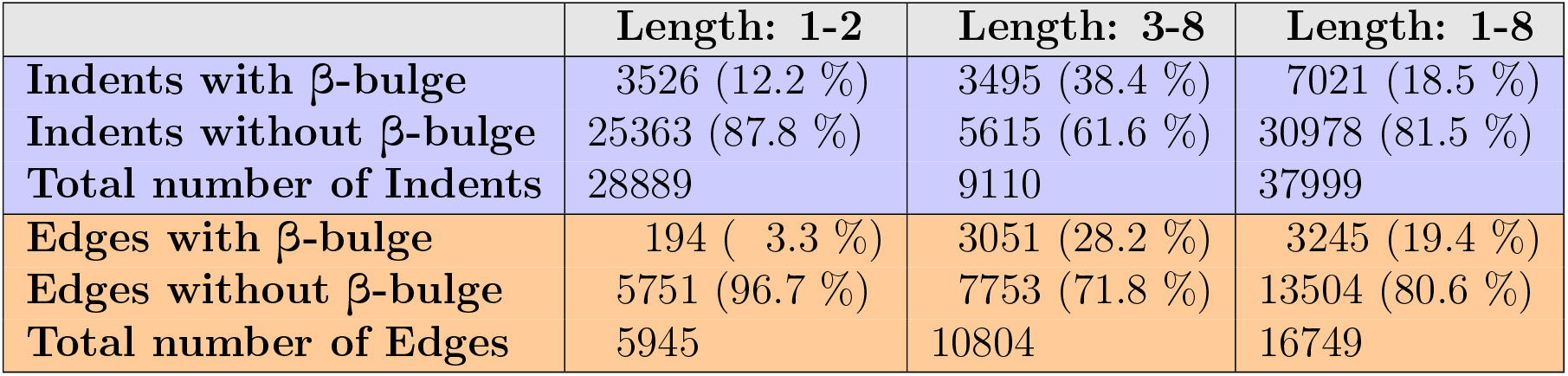
Strand statistics

|  | Length: 1-2 | Length: 3-8 | Length: 1-8 |
| --- | --- | --- | --- |
| Indents with β-bulge | 3526 (12.2 %) | 3495 (38.4 %) | 7021 (18.5 %) |
| Indents without β-bulge | 25363 (87.8 %) | 5615 (61.6 %) | 30978 (81.5 %) |
| Total number of Indents | 28889 | 9110 | 37999 |
| Edges with β-bulge | 194 ( 3.3 %) | 3051 (28.2 %) | 3245 (19.4 %) |
| Edges without β-bulge | 5751 (96.7 %) | 7753 (71.8 %) | 13504 (80.6 %) |
| Total number of Edges | 5945 | 10804 | 16749 |

Very short β-strands (lengths 1 or 2 residues) are naturally less suitable for β-sheet propagation by stacking of further β-strands. Properties of such short β-strands are expected to be different from longer β-strands. Therefore, β-strands were grouped in two categories, viz. short (1-2 amino acid residues) and long (3-8 amino acid residues). The distribution of indents and edges changed noticeably when these two categories were considered. Approximately 76 % of the indents are short indents whereas only 36 % of the edges fall into the category of short edges. Thus, indents tend to be shorter while longer edges are more common. While the abundance of shorter indents naturally protects β-sheets, the longer indents do exist and it will be interesting to see if longer indents incorporate different amino acid residues as a strategy to arrest β-sheet propagation. We will discuss this aspect further in this paper.

### Comparison of indents and edges with helices, surface residues and disordered residues

The effects of different localization of indents and edges could be reflected in their amino acid residue composition. Therefore, we studied the enrichment and depletion of various amino acid residues in different types of β-strands using Composition profiler [30]. Helices (α, 3_10_, π) form a major class of secondary structures that markedly differ from β-strands and β-sheets. Comparing β-strands with helices revealed mostly similar amino acid enrichments and depletions in both indents and edges (Table 2).

**Table 2:**
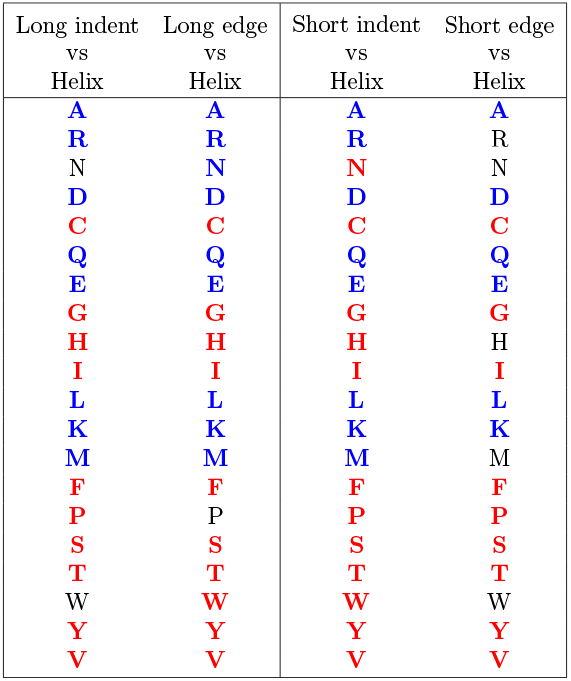
Comparisons of indents and edges with helices (α, 3_10_, π) Red: Enrichment (P-value ≤ 0.05), Blue: Depletion (P-value ≤ 0.05), Black: Not significant (P-value > 0.05) [30]

In general indents and edges are both enriched in hydrophobic and bulky residues compared to helices. Nonetheless, there are enrichments of small residues including serine, theronine, proline and glycine, that might be involved in anti-stacking strategies of β-strands.

When indents and edges were compared with surface residues (as defined by Vacic et al. [30]), both edges and indents showed similar enrichments of bulky residues (Table 5). The observed dissimilarity with the surface residues indicates that indents as well as edges, despite being on the periphery of β-sheet, are not as prone to solvent exposure as the known surface residues.

**Table 3:**
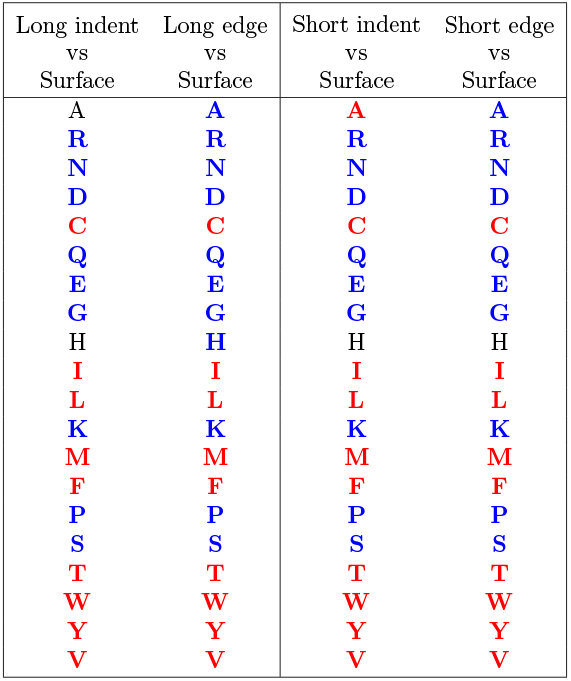
Comparisons of indents and edges with surface residues. Red: Enrichment (P-value ≤ 0.05), Blue: Depletion (P-value ≤ 0.05), Black: Not significant (P-value > 0.05), Surface: Surface residues determined by the Molecular Surface Package [30]

Indents and edges were also compared with a set of known disordered residues obtained from the Disprot database [31]. Compared to the Disprot data, indents and edges seem to be depleted in disorder promoting residues such as proline, glycine, aspactic acid, glutamic acid and glutamine.

Enrichment and depletion of various amino acid residues with respect to helices, surface residues and disordered residues fails to distinguish between indents and edges. This is expected because the amino acid preferences for helices, surface residues and disordered residues are very different from those for β-strands of any kind, thereby masking the difference between the subclasses of β-strands. Therefore, direct comparisons of composition profiles of indents and edges were performed.

**Table 4:**
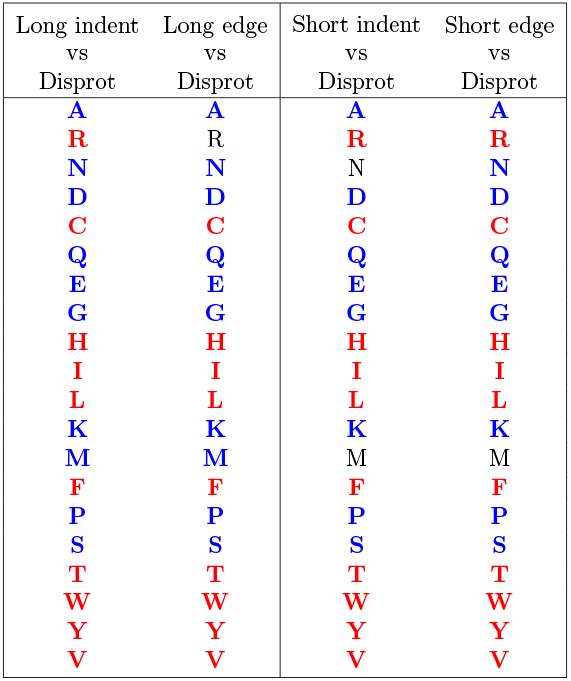
Comparisons of indents and edges with disordered residues. Red: Enrichment (P-value ≤ 0.05), Blue: Depletion (P-value ≤ 0.05), Black: Not significant (P-value > 0.05), Disprot: Set of disordered residues from Disprot 3.4 [30]

### Comparison of indents with edges

Solvent accessible surface area is an important feature that one can expect to be a point of difference between indents and edges due to their positioning in β-sheets. Indeed, the relative solvent accessibility of residues in indents and edges shows two distinct peaks as shown in the histogram, Figure 3.

**Figure 3:**
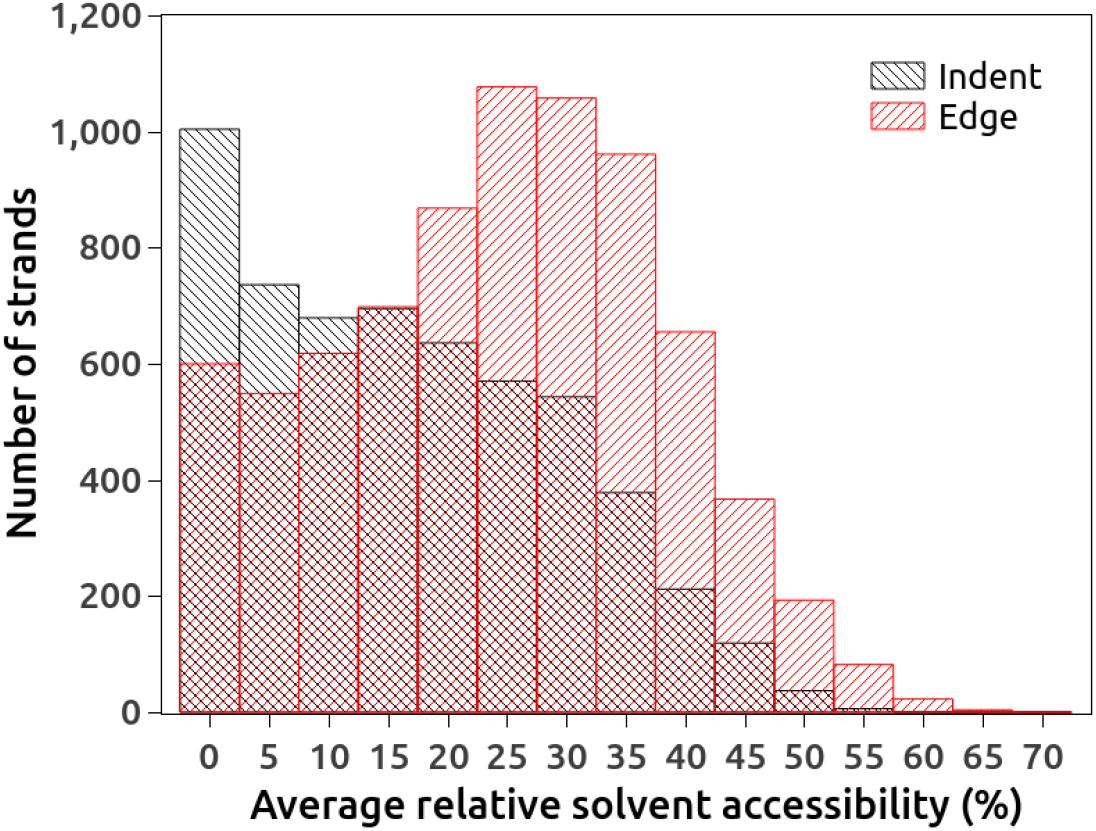
Solvent accessibility of long indents and long edges

Amino acid composition is another aspect based on which indents and edges can be compared. As shown in the table 5, long indents show enrichment of alanine, asparagine, aspartic acid, glycine and proline residues compared to long edges. Short indents also show enrichment of similar amino acid residues, except for a noticeable depletion in proline content compared to short edges. The long indents are also depleted in bulky/hydrophobic residues such as phenylalanine, tryptophan, tyrosine, isoleucine and valine, which is not true in case of comparison between short indents and edges.

**Table 5:**
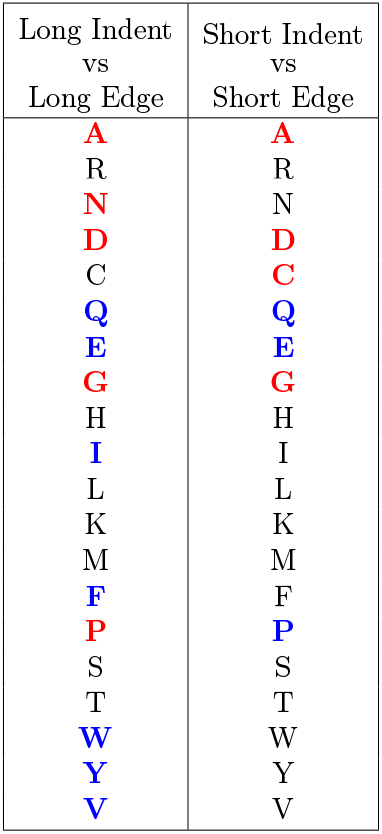
Comparisons of indents and edges with disordered residues. Red: Enrichment (P-value ≤ 0.05), Blue: Depletion (P-value ≤ 0.05), Black: Not significant (P-value > 0.05), [30]

Indents and edges were also found to differ from each other with respect to β-bulges. β-bulges form irregularities in periodic main-chain hydrogen bonds between adjacent β-strands. Work done by Richardson et al. [32] and Chan et al. [33] forms basis for the definition of β-bulges. Chan et al. [33] have systematically classified β-bulges. A revised analysis by Craveur et al. [34] describes how β-bulges impart irregularities in β-strand pairing by disrupting alternation of side-chains or terminate the β-strand. Our analysis showed that approximately 18-20 % of indents as well as edges have β-bulges (Table 1). Figure 4 shows relative frequencies of indents and edges for different strand lengths. The overall length distribution profiles of indents and edges does not show much difference. However, when the population is segregated based on occurrence of one or more β-bulges, the differences become visible. Indents and edges containing β-bulges show opposite trends in the strand length distribution (Figure 4 - top right). The short indents with β-bulges occur much more frequently than the long indents, whereas, long edges containing β-bulges are more common than the short edges. This shows that the longer edges might be using β-bulges as a protection against aggregation by favoring β-sheet termination. The longer indents, on the other hand, might get protected by various other mechanisms.

**Figure 4:**
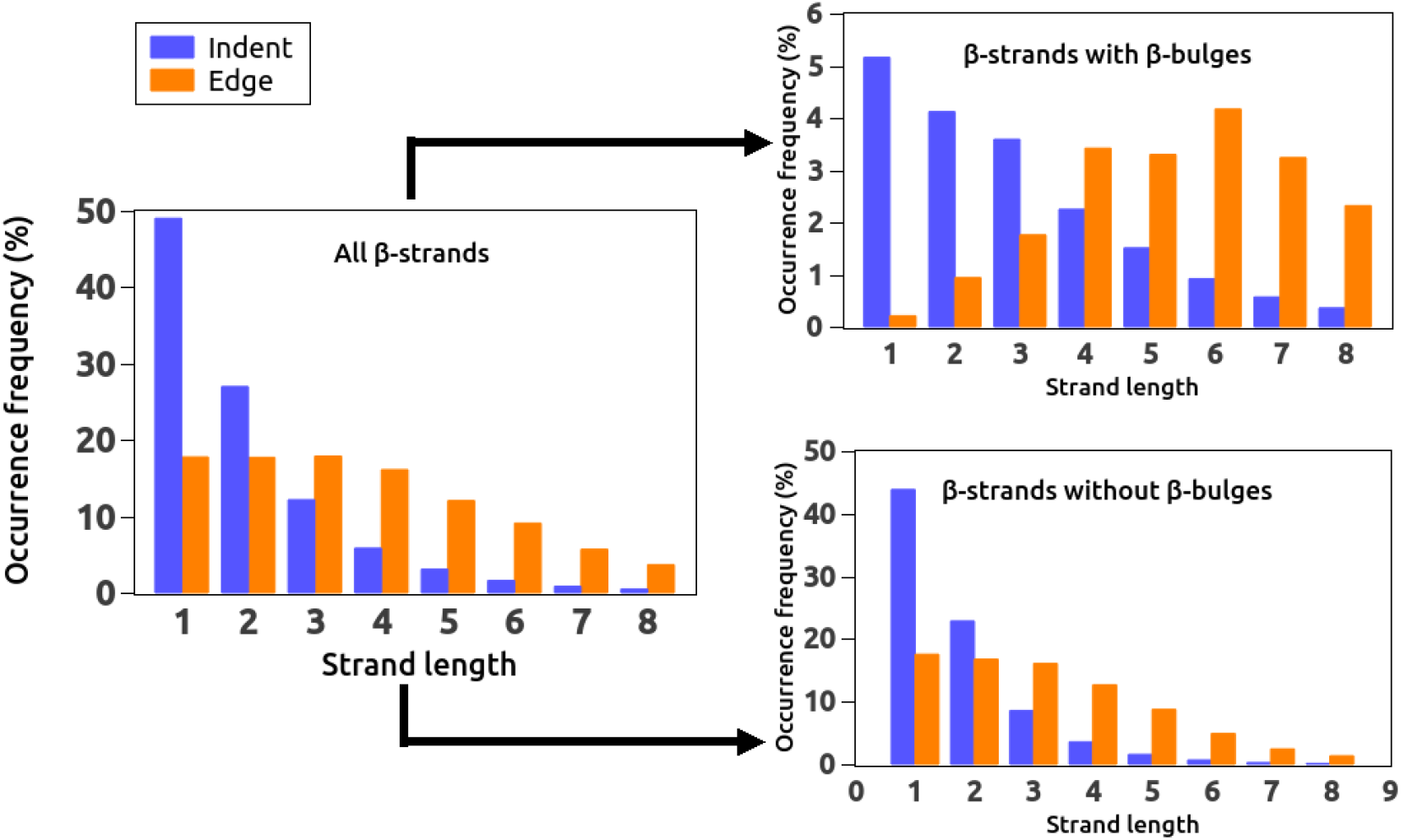
Length distribution, Left: All indents and edges, Top right: Indents and edges with β-bulges, Bottom right: Indents and edges without β-bulges

### Comparison of long strands with short strands

Length of a β-strand is a very important factor that governs stacking of further β-strands. Longer edges as well as indents are prone to further stacking and are known to possess various strategies to avoid strand pairing [19]. The short edges and indents, on the other hand, are inherently protected from stacking of other β-strands. To get an idea about which strategies are present in edges and indents, longer strands were compared with shorter strands of the same type. This comparison provided information on which other amino acid residues get incorporated in longer strands to make them resistant to further β-strand stacking. The amino acid residues and properties that are enriched in the longer strands in comparison with the shorter strands indicated the strategies used by the longer strands to avoid β-sheet propagation.

Table 6 shows comparison of long and short β-strands (indents as well as edges). The long indents seem to be enriched in glutamine, glutamic acid, proline and valine compared to short indents, while long edges show enrichment of glutamic acid, isoleucine and valine compared to short edges (Table 6a). As opposed to the comparison between long and short indents, proline is depleted in long edges compared to short edges. To better understand these observations, the comparisons were made in terms of various properties as shown in table 6b. The long indents seem to be less hydrophobic and enriched in flexible residues with higher solvation potential compared to the short indents. Edges show the opposite trend with the long edges being more hydrophobic than the short edges.

**Table 6.**
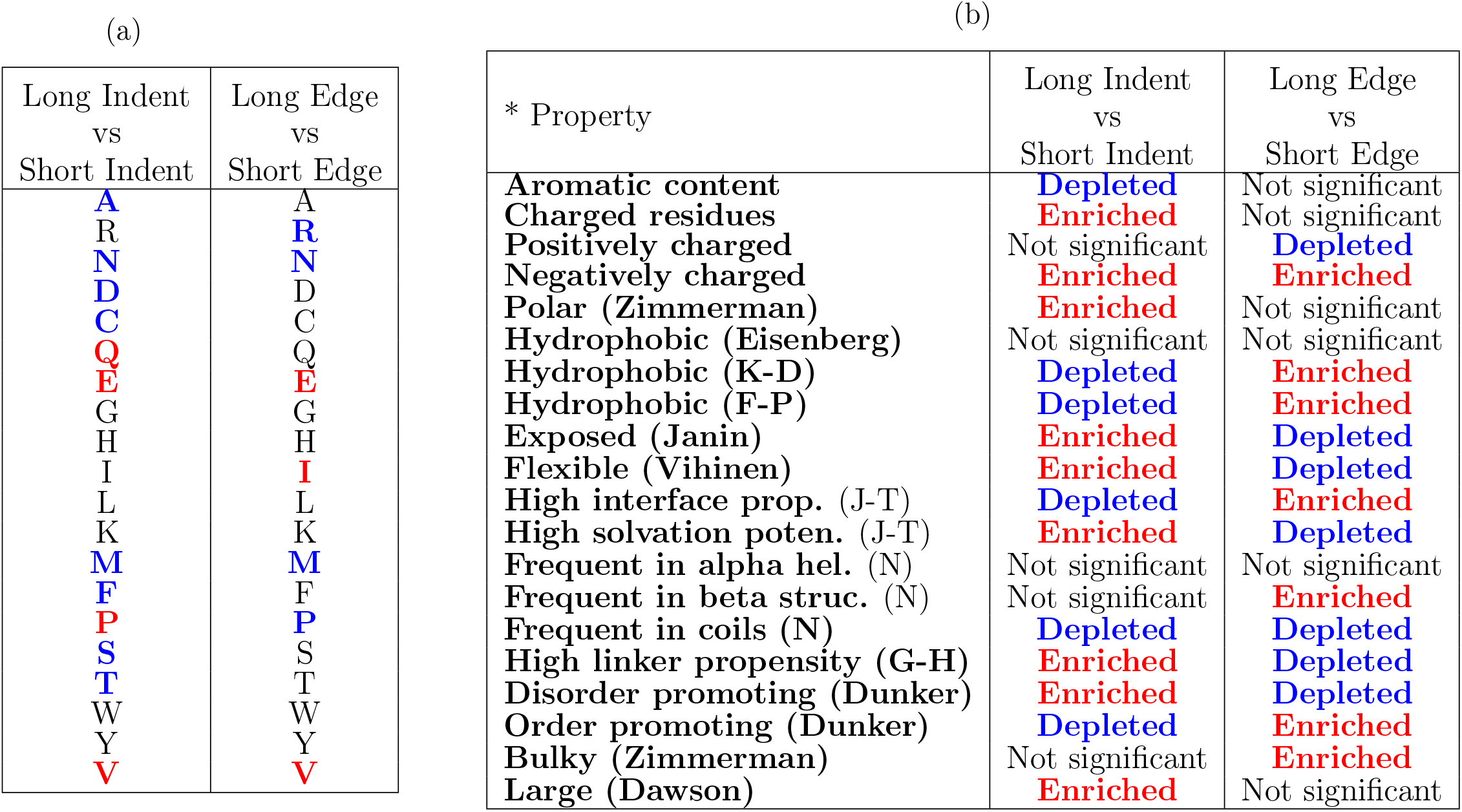
Enrichment, and depletion of (a) Amino acid residues, (b) Properties. Red: Enrichment (P-value ≤ 0.05), Blue: Depletion (P-value ≤ 0.05), Black: Not significant (P-value > 0.05). Definitions of the properties can be found in Vacic et al. [30].

These observations can be explained as follows. The indents and edges share a common feature i.e. lack of β-bridge partner on one side, thereby rendering the backbone available for hydrogen bonding. However, indents and edges occupy different regions of the β-sheet which is also reflected to some extent in their localization within the folded protein. The localization within the folded protein molecule affects the strategies used by the indents and edges to avoid β-sheet propagation.

The indents are in general less solvent exposed (Figure 3). The buried nature of indents raises the risk of β-strand stacking with increasing length. Therefore, it seems justified for long indents to be less hydrophobic. The shorter indents are already protected from further β-strand interactions due to their reduced length.

In contrast to indents, edges are generally more solvent exposed (Figure 3). Therefore, the protein fold has already taken care of avoiding further β-strand stacking, to some extent, from within the same protein molecule. However, long edges still need to be protected from intermolecular β-strand stacking. Figure 4 (top right) shows that longer edges are frequently involved inβ-bulges which is known to be a negative design principle to avoid β-strand stacking [19].

## Conclusion

Indents and edges form the periphery of β-sheets. Our work shows that indents are quite common in β-sheets. On an average there is at least one indent in each inner strand. Our analysis identifies subtle differences between properties of indents and edges. Similar to edge strands, indents also provide protection from unhindered β-sheet propagation. However, unlike edge strands, indents are extended edge-like portions of inner strands of β-sheets, thus occupying different position within β-sheets compared to edge strands. Therefore, understanding the differences between indents and edges might be useful for *de novo* design of β-sheets. Based on the present analysis, following features of indents and edges could be highlighted.

- Short indents are more common than short edges. Approximately 76 % of the indents are short indents while ≈36 % edges are short edges.
- Indents show lower proportion of β-bulges with increasing length, whereas, edges show higher proportion of β-bulges with increasing length.
- Edges are generally more solvent exposed than the indents.
- Long indents are at higher risk of aggregation due to their mostly buried nature and therefore are depleted in hydrophobic residues with an enrichment of flexible residues with high solvation potential as a strategy to deter further β-strand stacking.
- Long edges seem to better tolerate hydrophobic and bulky residues, however, they are known to be protected from β-strand stacking through mechanisms such as strategically placed charges, loop coverage and inward-pointing charged side chains.

## Acknowledgement

The authors thank the Indian Council of Medical Research (ICMR) for fellowship to HK.

